# Historical contingency in the evolution of antibiotic resistance after decades of relaxed selection

**DOI:** 10.1101/695767

**Authors:** Kyle J. Card, Thomas LaBar, Jasper B. Gomez, Richard E. Lenski

## Abstract

Populations often encounter changed environments that remove selection for the maintenance of particular phenotypic traits. The resulting genetic decay of those traits under relaxed selection reduces an organism’s fitness in its prior environment. However, whether and how such decay alters the subsequent evolvability of a population upon restoration of selection for a previously diminished trait is not well understood. We addressed this question using *Escherichia coli* strains from the long-term evolution experiment (LTEE) that independently evolved for multiple decades in the absence of antibiotics. We first confirmed that these derived strains are typically more sensitive to various antibiotics than their common ancestor. We then subjected the ancestral and derived strains to various concentrations of these drugs to examine their potential to evolve increased resistance. We found that evolvability was idiosyncratic with respect to initial genotype; that is, the derived strains did not generally compensate for their greater susceptibility by “catching up” to the resistance level of the ancestor. Instead, the capacity to evolve increased resistance was constrained in some backgrounds, implying that evolvability depended upon prior mutations in a historically contingent fashion. We further subjected a time-series of clones from one LTEE population to tetracycline and determined that an evolutionary constraint arose early in that population, corroborating the role of contingency. In summary, relaxed selection not only can drive populations to increased antibiotic susceptibility, but it can also affect the subsequent evolvability of antibiotic resistance in an unpredictable manner. This conclusion has potential implications for public health, and it underscores the need to consider the genetic context of pathogens when designing drug-treatment strategies.

## Introduction

A population may encounter an environmental change that removes or reduces a selective pressure that was previously important for the maintenance of a trait (Darwin 1859; Lahti et al. 2009). Adaptation to the new environment can therefore affect an organism’s fitness in its prior environment. These correlated responses may lead to the functional decay of unused traits over time or, conversely, their maintenance despite relaxed selection (Lahti et al. 2009). However, the evolutionary processes driving these responses are often hard to disentangle because one must rely on retrospective studies and historical inference.

By contrast, evolution experiments with microorganisms provide a powerful approach to study correlated responses. Microbes often have large population sizes and fast generations, and they are amenable to freezing and revival. One can therefore observe evolution in action, directly compare ancestral and derived forms, and simultaneously assess adaptation to one environment and quantify correlated fitness responses in another. Accordingly, numerous studies with bacteria (Chao et al. 1977; Lenski 1988; Reboud & Bell 1997; Cooper & Lenski 2000; Cooper et al. 2001; Ellis & Cooper 2010; Leiby & Marx 2014), viruses (Turner and Elena 2000; Duffy et al. 2006; Agudelo-Romero et al. 2008; Coffey and Vignuzzi 2011; Wasik et al. 2015; Meyer et al. 2016), and yeast (Wenger et al. 2011; Ratcliff et al. 2012; Koschwanez et al. 2013) have found that fitness tradeoffs between environments are common.

Tradeoffs are often caused by antagonistic pleiotropy, which occurs when a mutation that is beneficial in one environment is deleterious in another. This process can have important public-health consequences when antibiotic-resistance mutations or acquired resistance genes impose costs on bacterial growth and competitiveness relative to their sensitive counterparts in the absence of drugs (Lenski 1997; Andersson and Hughes 2010). Previous studies have shown that pleiotropic fitness costs are widespread among resistance determinants to diverse drug classes (Nguyen et al. 1989; Schrag et al. 1997; Rozen et al. 2007; Han et al. 2009), although their magnitudes are variable and may also depend on the genetic background (Lenski et al. 1994; Andersson and Hughes 2010; Melnyk et al. 2015; Palmer et al. 2018)

Given that antibiotic-resistance mutations and genes commonly impose fitness costs, one would expect that resistance should decline over time in the absence of antibiotic exposure. However, compensatory evolution often reduces or eliminates these tradeoffs (Bouma and Lenski 1988; Schrag et al. 1997; Reynolds 2000; Rozen et al. 2007). Adaptive trends during compensatory evolution have been studied using a number of *Escherichia coli* mutants resistant to the drug rifampicin (Barrick et al. 2010). That study found that the mutants were generally less fit than their sensitive progenitors in a permissive antibiotic-free environment; moreover, the compensatory effects of subsequent beneficial mutations were greater when the resistance was more costly. Thus, compensation exhibited a pattern of diminishing-returns adaptation in that study.

Even when bacteria have no known history of exposure to antibiotics, they may have low-level resistance to some drugs because of intrinsic structural or functional features, including their cell envelope and efflux pumps (Cox and Wright 2013). However, even intrinsic resistance may decline in the absence of drug exposure, if relevant genes accumulate mutations either by selection or drift in permissive environments (Cooper and Lenski 2000).

A recent study used the *E. coli* long-term evolution experiment (LTEE), and antibiotic resistance as a model trait, to study changes in an organism’s capacity to tolerate environmental stresses when it evolves for a long period in the absence of those stresses (Lamrabet et al. 2019). In the LTEE, twelve replicate populations were founded from a common ancestor and have been propagated daily for over 30 years in a medium without antibiotics (Lenski et al. 1991; Tenaillon et al. 2016). In particular, Lamrabet et al. measured changes in mostly low-level intrinsic resistance between ancestral and derived strains isolated from each population after generations 2,000 and 50,000. They found that derived strains were usually more susceptible to most antibiotics than their ancestor, and from multiple lines of evidence they inferred that these losses of intrinsic resistance resulted primarily from pleiotropic side-effects of beneficial mutations that arose during the LTEE.

Taken together, the experimental evolution studies described above have two contrasting implications relevant for medicine and public health. First, resistance to antibiotics (including even low-level intrinsic resistance) may decline in the absence of drug exposure. Second, evolution can often compensate for deleterious side-effects of mutations, thereby facilitating the maintenance of evolved resistance. The question then arises how readily bacteria can overcome losses of antibiotic resistance that arose during periods of relaxed selection through subsequent evolution in the presence of drugs. In this study, we address this fundamental question by using the LTEE ancestor and derived strains isolated from four populations after 50,000-generations to examine how evolution in the absence of antibiotics affects the bacteria’s potential to evolve increased resistance when drugs are introduced. In so doing, we examine the role that genetic background plays in resistance evolvability (Figure 1). Does resistance evolution tend to follow a general trend of diminishing returns (Barrick et al. 2010; Wiser et al. 2013; Kryazhimskiy et al. 2014), such that derived strains that are initially more susceptible to a drug can increase their resistance disproportionately relative to their ancestor (Figure 1b)? Or is evolvability idiosyncratic with respect to prior evolutionary history (Blount et al. 2008, 2018), such that the relative gains in resistance are independent of a strain’s initial susceptibility (Figure 1c)?

**Figure 1.**
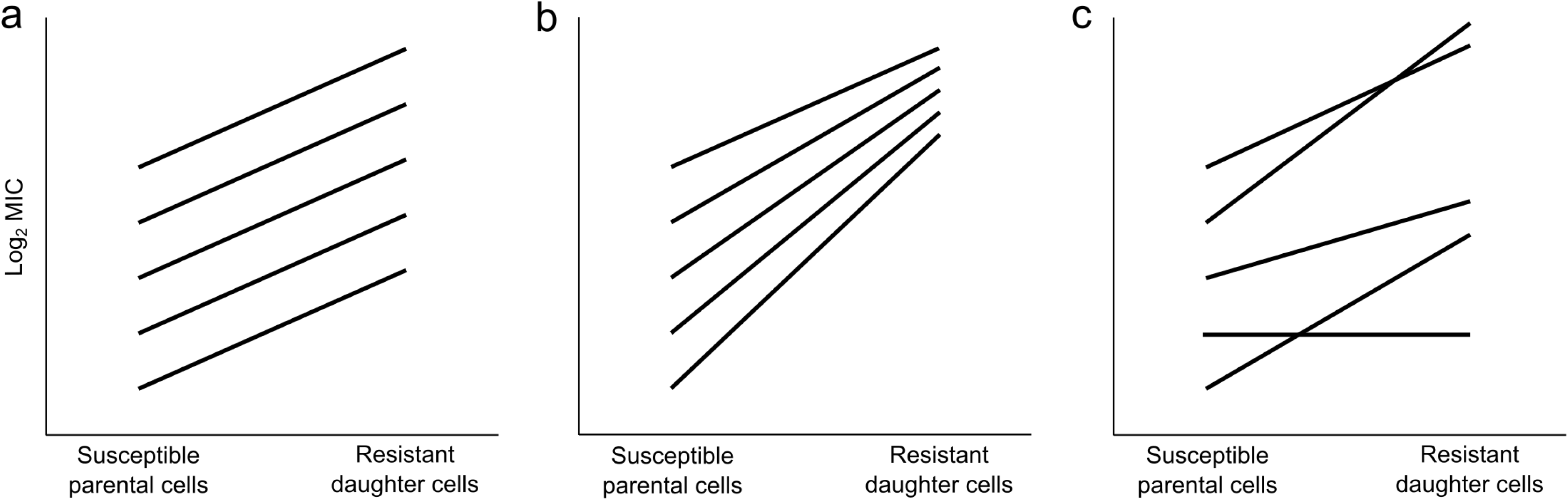
Schematic illustration of the evolvability of antibiotic resistance under three scenarios. A strain’s evolvability is defined operationally as the maximum increase in resistance from an initially susceptible genotype during one round of drug selection. (**a**) Null model, with no effect of genetic background on evolvability. (**b**) Diminishing-returns model, such that backgrounds with low initial resistance are more evolvable than backgrounds that are initially more resistant. (**c**) Idiosyncratic-effects model, in which evolvability varies among genetic backgrounds but is uncorrelated with their initial level of resistance.

We confirmed the finding of Lambaret et al. (2019) that the LTEE-derived strains had typically become more susceptible to antibiotics during relaxed selection. However, contrary to our expectation based on a diminishing-returns model, we discovered that these derived strains were usually no more evolvable (and sometimes less evolvable) than their ancestor when exposed to various antibiotics. Instead, idiosyncratic responses dominated over any diminishing-returns tendency, such that the capacity to evolve resistance was hampered on some LTEE-derived genetic backgrounds. These results indicate that evolution and diversification of a single bacterial species in a permissive environment can lead to unpredictable changes in their potential to evolve antibiotic resistance. Our work suggests that improved methods for predicting, at the strain level, a pathogen’s evolutionary potential should become an integral component of resistance surveillance and patient treatment in light of the global threat of antibiotic resistance.

## Materials and Methods

### Bacterial strains

All of the strains used in this study are from the *E. coli* long-term evolution experiment (LTEE). In the LTEE, twelve replicate populations were founded from a common ancestral strain, called REL606 (Lenski et al. 1991). These populations have been propagated for over 31 years by daily 1:100 transfers in glucose-supplemented Davis minimal (DM) medium without any antibiotics (Lenski et al. 1991), resulting in >71,000 cell generations to date. Samples from each population are frozen periodically at –80°C. In this study, we quantified the intrinsic antibiotic resistance and evolvability of the ancestral and derived clones isolated from four populations (designated Ara–5, Ara–6, Ara+4, and Ara+5) after 50,000 generations of the LTEE. We chose these strains for two reasons. First, the source populations of these derived clones retained the low ancestral mutation rate, and therefore they accumulated many fewer mutations than their counterparts from several populations that evolved hypermutability (Tenaillon et al. 2016). This characteristic should increase the tractability of identifying candidate alleles affecting resistance evolvability, which we hope to achieve in future work. Second, generation 50,000 is the latest generation for which whole-genome sequence data are available for the clonal samples (Tenaillon et al. 2016).

We also examined when the Ara+5 population evolved a diminished capacity to increase its tetracycline resistance (as described in the Results) by testing two strains isolated from this population at several earlier time points (generations 500, 1,000, 1,500, 2,000, 5,000, and 10,000). All of the strains used in this study are listed in Table 1.

**Table 1.** Bacterial strains used in this study.

| **Strain** | **Generation** | **LTEE Population** |
| --- | --- | --- |
| REL606 | 0 | - |
| REL772A | 500 | Ara+5 |
| REL772B | 500 | Ara+5 |
| REL962A | 1,000 | Ara+5 |
| REL962B | 1,000 | Ara+5 |
| REL1066A | 1,500 | Ara+5 |
| REL1066B | 1,500 | Ara+5 |
| REL1162A | 2,000 | Ara+5 |
| REL1162B | 2,000 | Ara+5 |
| REL2177A | 5,000 | Ara+5 |
| REL2177B | 5,000 | Ara+5 |
| REL4534A | 10,000 | Ara+5 |
| REL4534B | 10,000 | Ara+5 |
| REL11339 | 50,000 | Ara-5 |
| REL11389 | 50,000 | Ara-6 |
| REL11348 | 50,000 | Ara+4 |
| REL11367 | 50,000 | Ara+5 |

All clones were derived from REL606, the ancestral strain of the LTEE (Lenski et al. 1991).

### Culture conditions and measurements of resistance and evolvability

All experiments were performed at 37°C. Bacterial strains were revived from frozen stocks by overnight growth in Luria Bertani (LB) medium. Cells from these cultures were then streaked onto DM agar plates supplemented with 4 mg/mL glucose. We randomly picked single isolated colonies from these plates to start multiple replicate populations in LB. Final population sizes in the LB cultures were approximately 1 – 2 × 10^9^ cells/mL. When an initially susceptible cell expands into a colony and then a population, new mutations spontaneously occur and increase in number during growth (Luria and Delbrück 1943). The evolution of antibiotic-resistant mutants will therefore originate by independent mutational events in each replicate population (Luria and Delbrück 1943; Kassen and Bataillon 2006).

We define a strain’s evolvability as the maximum increase in antibiotic resistance from an initially susceptible genotype during one round of drug selection. Evolvability experiments were performed using Mueller-Hinton (MH) agar (Acumedia) supplemented with 1 mg/mL glucose, 0.1 mg/mL magnesium sulfate, 0.01 mg/mL thiamine, and a series of two-fold dilutions of an antibiotic. We chose four antibiotics to use in our study because they have diverse cellular targets: ampicillin and ceftriaxone inhibit cell-wall synthesis, ciprofloxacin inhibits DNA replication, and tetracycline inhibits protein synthesis. We prepared stock solutions of each antibiotic following the manufacturers’ instructions, which were then stored at –20°C.

One-mL samples of each population were centrifuged at 8,000 rpm for 2 min and resuspended in an equal volume of saline. We then plated 100 µL (containing ∼1 – 2 × 10^8^ cells) from each suspension onto the antibiotic-amended MH agar plates, and minimum inhibitory concentrations (MICs) were evaluated after 48 h of incubation. For this study, we operationally define a pair of MICs for each series of antibiotic-amended plates as the lowest concentration that prevents either confluent growth or isolated colonies. According to this approach, confluence indicates growth by the susceptible “parental” strain, while isolated colonies are resistant “daughter” mutants. A strain’s evolvability was calculated from the difference in MIC between these two genotypes. Putative resistant mutants were confirmed by streaking them onto fresh antibiotic-amended MH plates. This approach ensured that a selected clone was indeed a resistant mutant with a stably inherited increase in its MIC, as opposed to a so-called “persister” that exhibited higher-than-average phenotypic tolerance relative to genetically identical cells (Balaban et al. 2019). Cultures of mutant clones were then frozen at –80°C in LB medium supplemented with 15% glycerol as a cryoprotectant.

Figure 2 provides a schematic representation of our methods for measuring the MICs of sensitive parental strains and their resistant daughter derivatives. In Figure 3, we show images of the resulting plates for one replicate series across a 256-fold (= 2^8^) range of ciprofloxacin concentrations for the LTEE ancestral clone. In these images, one sees confluent growth on the first 3 plates, isolated colonies on the next 2 plates, and no evident growth on the 4 plates with the highest concentrations. Based on these plates, we scored the MIC of the sensitive parental strain as the lowest concentration that inhibited confluent growth, which was 0.0025 µg/mL in this example. We scored the MIC of the resistant daughter derivative as the lowest concentration where even isolated colonies were absent, in this case 0.01 µg/mL. The log_2_-transformed difference between these values (i.e., log_2_ 0.01/0.0025 = 2 in this example) provides one estimate of the evolvability of the LTEE ancestral strain with respect to ciprofloxacin. We obtained 32 independent estimates of these MICs and the associated evolvabilities for the ancestral strain against each of the four antibiotics used in our study. We similarly obtained eight independent estimates of the MICs and associated evolvabilities for each of the four 50,000-generation strains used in our study against each of the same antibiotics. Photographs of all of the replicate plate series used to estimate these values have been archived on the Dryad Digital Repository (doi pending publication).

**Figure 2.**
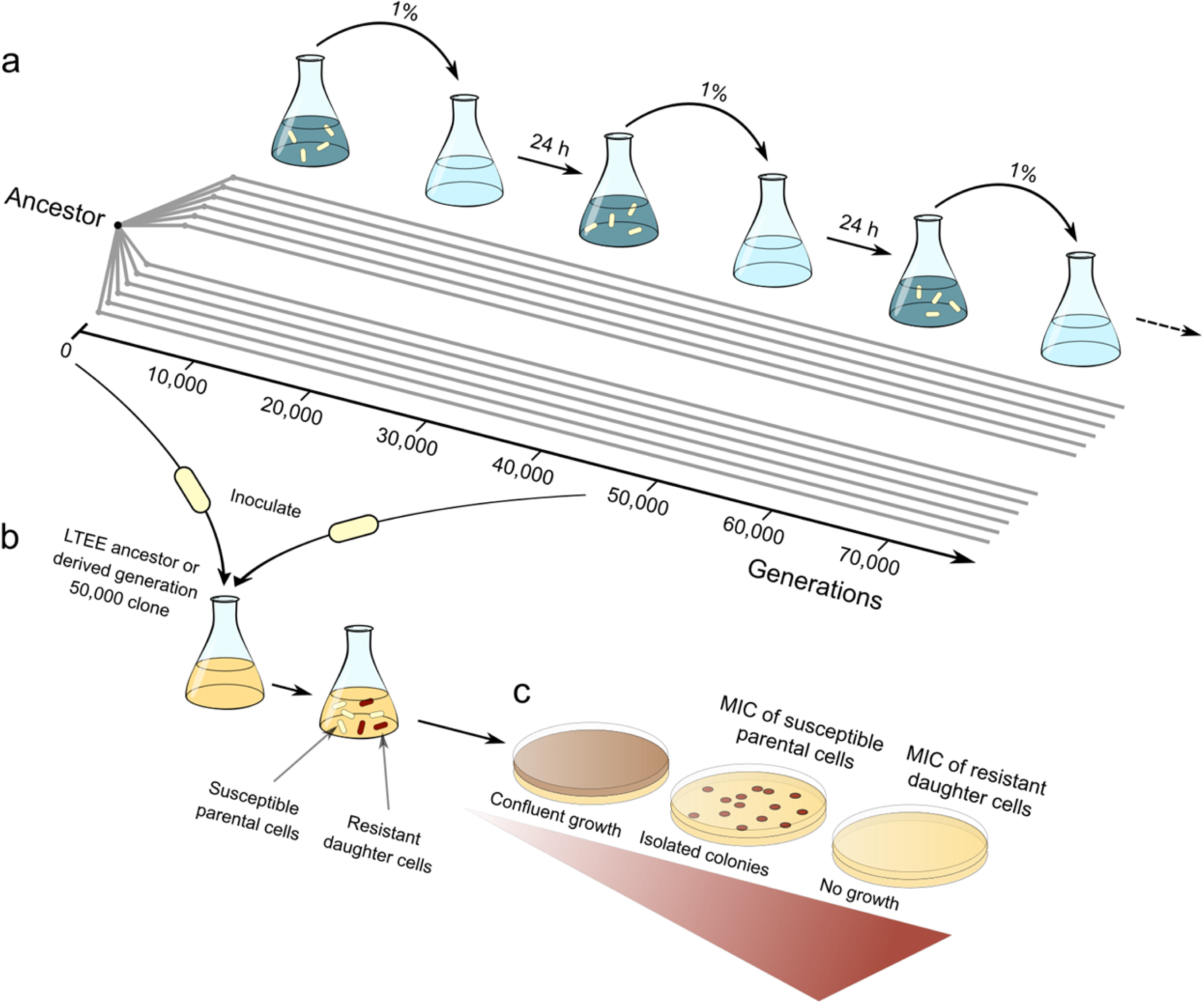
Schematic illustration of the long-term evolution experiment (LTEE) and evolvability study design. (**a**) Twelve initially identical *E. coli* populations were founded from a common ancestor to start the LTEE. These populations have evolved for >71,000 generations with daily serial transfers in a growth medium without antibiotics. (**b**) In this study, antibiotic-susceptible ancestral or derived clones from generation 50,000 were inoculated into replicate cultures. A resistance mutation may arise spontaneously and increase in number during a population’s expansion, resulting in two genetic variants: the susceptible parental cells and their descendent resistant daughters. (**c**) These whole populations are then spread onto agar plates supplemented with two-fold increasing concentrations of an antibiotic (shown in red). Minimum inhibitory concentrations (MICs) of these two variants correspond to the lowest antibiotic concentration that inhibits confluent growth and that prevents even isolated colonies, respectively. Resistant clones are confirmed by streaking onto fresh plates with relevant antibiotic concentrations.

**Figure 3.**
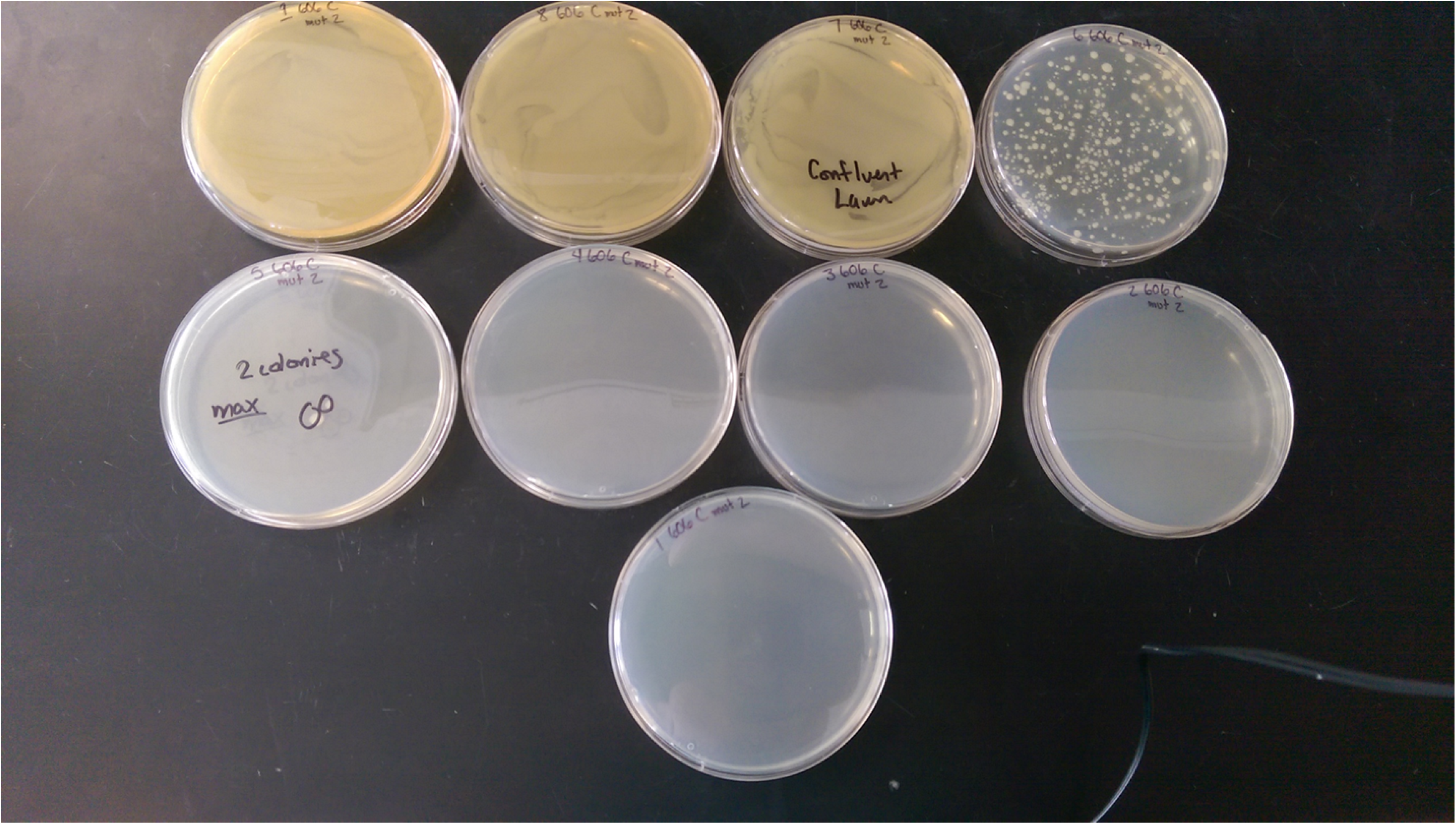
Experimental plates. Whole populations containing susceptible parental and resistant daughter cells were spread onto MH agar amended with two-fold increasing concentrations of an antibiotic (left to right, and down). Confluent lawns of bacterial growth (plates 1–3) consist largely of drug-susceptible cells. Isolated colonies (plates 4–5) are putatively resistant mutants. See Supplementary Data for images of all experimental plates.

### Experimental design and data analyses

All MIC values were transformed by taking their base-2 logarithm because the antibiotic concentrations were tested across a series of two-fold dilutions. For each experimental block, an independently isolated LTEE ancestral clone was paired with each derived clone. We had two predictions when we began this study: (i) the derived bacteria would be more susceptible to antibiotics (lower MICs) than their common ancestor as a consequence of the relaxed selection they experienced in the permissive LTEE environment; and (ii) the derived bacteria would be more evolvable than their ancestor when challenged with antibiotics, following a general trend of diminishing-returns adaptation.

Statistical tests that rely on normally distributed data were deemed inappropriate for this study owing to the discrete, lumpy nature of the measurements. Instead, we used nonparametric methods. There were also numerous instances in which the derived clones were equal both in MIC and evolvability to the paired assays for the ancestor, and these ties introduced additional complications. Therefore, we used trinomial tests to examine changes in the direction of our expectations relative to the null hypotheses that changes are equally frequent in either direction (Bian et al. 2011). We performed these analyses by individually comparing the four derived clones with their paired ancestors across each antibiotic. Probabilities were then combined from these independent significance tests using Fisher’s method with 8 degrees of freedom (i.e., df = 2*k*; where *k* is the number of comparisons) (Fisher 1934; Sokal and Rohlf 1994). As explained previously (Fig. 1c), evolvability might be idiosyncratic and therefore not correlated with the initial level of resistance. To assess this possibility, we performed a Kruskal-Wallis one-way ANOVA to test for heterogeneity in evolvability among the LTEE lines. Datasets and R analysis scripts are available on the Dryad Digital Repository (doi pending publication).

## Results

### Antibiotic susceptibility profiles of the LTEE ancestral and derived clones

Antibiotic susceptibility measurements are generally quite repeatable (Figure 4). For each antibiotic, all 32 independent ancestral replicate MIC measurements were identical. Among the 16 sets of derived-clone replicates (4 clones × 4 antibiotics), the 8 replicate assays gave identical MICs in 2 cases (12.5%), they deviated minimally by a factor of 2 in 12 cases (75%), and in only 2 cases they deviated by a factor of 4 (12.5%). These results provide strong support for the use of our plate-based approach to quantify antibiotic susceptibility profiles.

**Figure 4.**
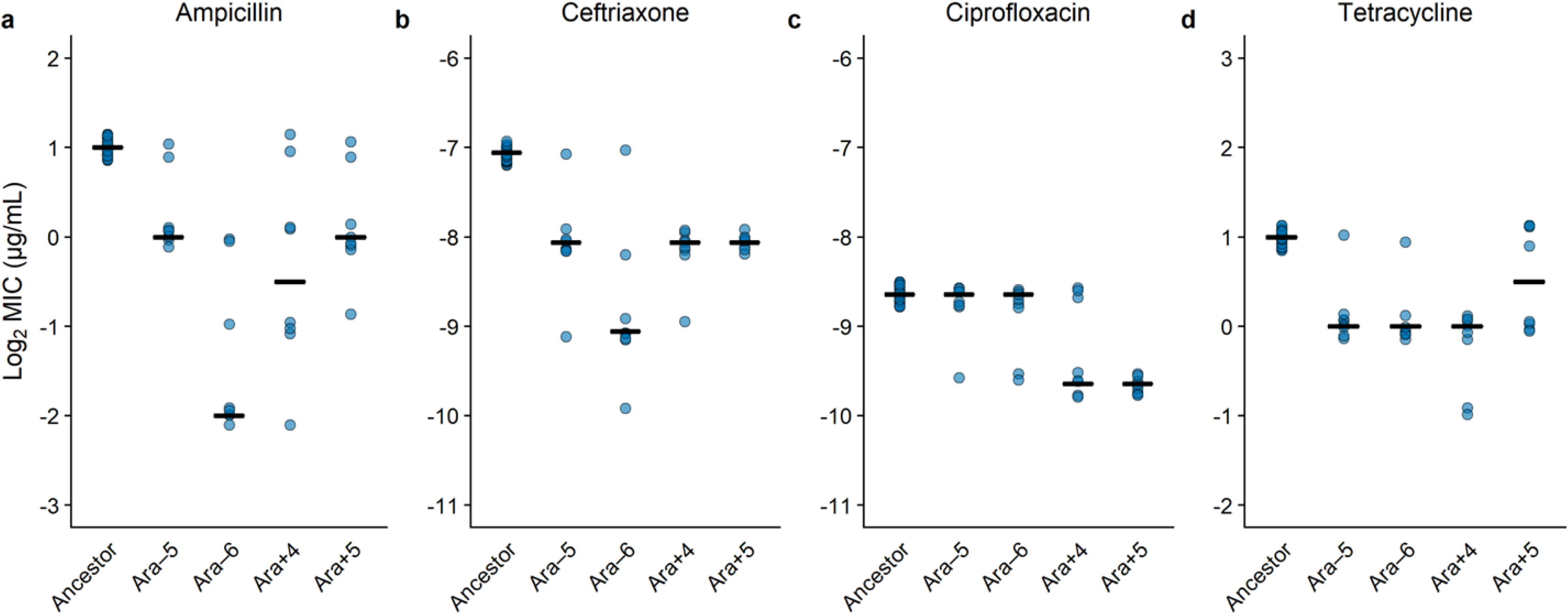
Intrinsic resistance usually declined over time in the absence of drug exposure. Comparison of the LTEE ancestor and four independently derived clones sampled after 50,000 generations for their susceptibilities to ampicillin, ceftriaxone, ciprofloxacin, and tetracycline (**a– d**). MICs are shown on a log_2_-transformed scale to reflect the fact that antibiotic concentrations were tested across a series of two-fold dilutions. In each panel, points show values obtained from 32 and 8 replicate assays for the ancestor and derived strains, respectively. Horizontal bars show the median of the log_2_-transformed MIC values for each strain on each antibiotic. The absolute values of the concentrations shown on the y-axis differ among the four antibiotics, but the range is the same in each panel.

### Changes in susceptibility under relaxed selection during the LTEE

For each antibiotic, we made 32 comparisons between the MICs of derived clones (4 clones × 8 replicates) against their paired and independently isolated ancestral clones. On balance, we observed increased susceptibility of the strains that evolved under relaxed selection (i.e., in the absence of antibiotics) during the LTEE, consistent with recently published results (Lamrabet et al. 2019) (Figure 4). All four derived strains have increased sensitivity to ampicillin (Fig. 4a), ceftriaxone (Fig. 4b), and tetracycline (Fig. 4d) relative to their common ancestor, and two of the derived strains were more susceptible to ciprofloxacin (Fig. 4c). These trends toward lower resistance are well supported by trinomial tests, as described in the Materials and Methods, and as shown in Tables 2 and S1.

**Table 2.** Statistical analyses of declines in intrinsic resistance during relaxed selection of clones sampled at generation 50,000 of the LTEE.

| **Antibiotic** | **$\chi^2$** | ***p*** |
| --- | --- | --- |
| Ampicillin | 40.06 | < 0.0001 |
| Ceftriaxone | 45.33 | < 0.0001 |
| Ciprofloxacin | 27.15 | 0.0007 |
| Tetracycline | 41.40 | < 0.0001 |

Analyses were performed using Fisher's combined probability method (df = 8) for multiple independent tests of the same hypothesis (Fisher 1934; Sokal and Rohlf 1994), with an underlying trinomial distribution for the null hypothesis (Bian et al. 2011).

### Evolvability profiles of the ancestor and derived clones

Next, we examined how the prior history of relaxed selection affected the evolvability of antibiotic resistance in the different genetic backgrounds. To address this question, we selected mutants of the ancestral and LTEE-derived strains that survived and grew sufficiently to form colonies at higher concentrations of the four antibiotics than their corresponding parental strains (Fig. 2). Recall that, in this study, we operationally define evolvability as the maximum observed increase in antibiotic resistance from an initially susceptible genotype during one round of drug selection (Fig. 1).

Evolvability measurements tended to be more variable than the MIC measurements. We examined the evolvability of 128 independent ancestral clones across the four antibiotics. There were 73 cases (57%) where these measurements corresponded to the median for that antibiotic, 49 cases (38.3%) where they differed by a factor of 2, and 6 cases (4.7%) where they differed by a factor of 4. Likewise, among the 16 sets of replicates for the LTEE-derived clones, the 8 assays varied by a factor of 2 in 12 cases (75%) and by a factor of 4 in 4 other cases (25%). The greater variation in evolvability measurements in comparison to MIC values among replicate assays is expected given the stochastic appearance of mutations in replicate cultures (Luria and Delbrück 1943), as well as the fact that increased resistance can occur through multiple mutational paths (Toprak et al. 2012; Baym et al. 2016).

### Effects of genetic background on the evolvability of resistance

We examined the possibility of two broad patterns of genetic-background effects with respect to resistance evolvability in our study. First, we asked whether evolvability followed a trend of diminishing returns, such that the more susceptible LTEE-derived genetic backgrounds generally produced mutants with proportionally greater gains in resistance than the ancestor. Both the ancestral and derived strains evolved resistance to varying degrees (Figure 5). The evolutionary potential of two of the four derived clones (Ara–5 and Ara–6) was noticeably greater relative to their ancestor in the ampicillin environment (Fig. 5a), but there were no clear instances of similar trends in the three other drug environments (Fig. 5b–d).

**Figure 5.**
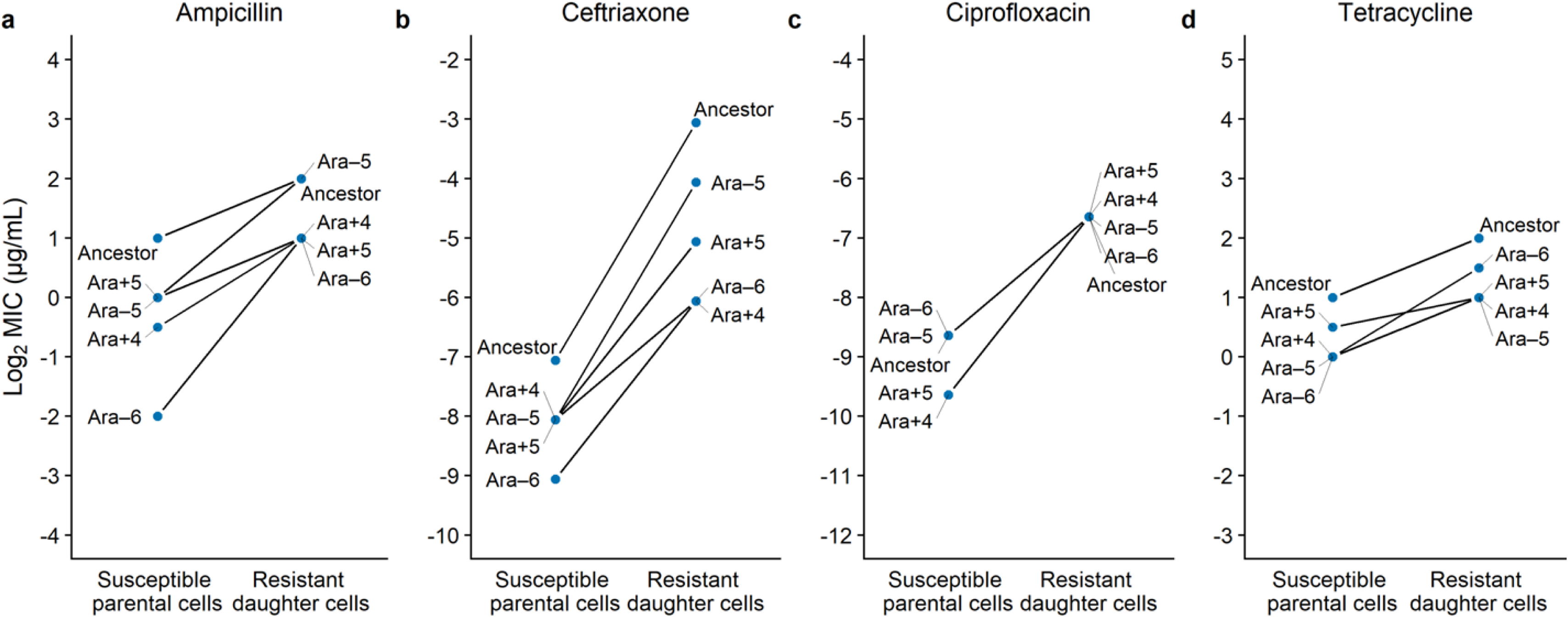
Genetic background affects the evolvability of LTEE lines exposed to antibiotics. Lines joining susceptible parental strains with their daughter mutants show the increases in resistance during one round of selection with the indicated antibiotic. If the slope of a derived strain is greater than that of the ancestor, then it has greater evolvability; and vice versa. Median MICs are shown on a log_2_-transformed scale to reflect the fact that antibiotic concentrations were tested across a series of two-fold dilutions. The y-axis ranges for the four drugs have been scaled to the ceftriaxone environment, which had the largest gains in resistance between the susceptible parental cells and the resistant daughter cells.

Overall, there was no statistical support for the diminishing-returns trend, despite the visual impression for the ampicillin treatment. We compared each derived strain’s evolutionary potential to its paired ancestor in the four drug environments. We used trinomial tests to quantify the likelihood that each derived strain’s evolvability was greater than its ancestral counterpart when tested against the null hypothesis of equally frequent changes in either direction, after taking into account the many numerical ties (Bian et al. 2011). Although the capacity of the derived Ara–5 clone to evolve increased resistance was significantly greater than its ancestor when considered in isolation (Table S2), it was marginally non-significant when we examined overall trends for each antibiotic (Table 3) using a meta-analysis approach (Fisher 1934; Sokal and Rohlf 1994).

**Table 3.** Statistical analyses of diminishing-returns trends in resistance evolvability of clones sampled at generation 50,000 of the LTEE.

| **Antibiotic** | **$\chi^2$** | ***p*** |
| --- | --- | --- |
| Ampicillin | 14.63 | 0.0668 |
| Ceftriaxone | 0.18 | 1 |
| Ciprofloxacin | 7.88 | 0.4456 |
| Tetracycline | 5.97 | 0.6511 |

Analyses were performed using Fisher's combined probability method ( $df = 8$ ) for multiple independent tests of the same hypothesis (Fisher 1934; Sokal and Rohlf 1994), with an underlying trinomial distribution for the null hypothesis (Bian et al. 2011).

We then asked whether the proportional resistance gains when exposed to the antibiotics were idiosyncratic among LTEE lines. For example, the capacity to evolve ceftriaxone resistance appeared to be reduced among three LTEE-derived backgrounds (Ara+5, Ara–6, and especially Ara+4) relative to their common ancestor (Fig. 5b). Similarly, the evolvability of the Ara+5 background with respect to tetracycline appears to be constrained (Fig. 5d). Indeed, this latter case was the only one in which the mutants of a strain systematically achieved a lower level of resistance than did the mutants of other strains that were initially more susceptible (indicated by the crossing lines in Fig. 5d). These idiosyncratic tendencies are statistically well supported by Kruskal-Wallis tests. For both ceftriaxone and tetracycline, these tests reject the null hypothesis of homogeneity in proportional resistance increases across the different genetic backgrounds (Table 4).

**Table 4.** Statistical analyses of idiosyncratic patterns in resistance evolvability of clones sampled at generation 50,000 of the LTEE.

| **Antibiotic** | **$\chi^2$** | ***p*** |
| --- | --- | --- |
| Ampicillin | 9.19 | 0.0566 |
| Ceftriaxone | 23.45 | 0.0001 |
| Ciprofloxacin | 7.59 | 0.1077 |
| Tetracycline | 10.18 | 0.0376 |

Analyses were performed using a Kruskal-Wallis one-way nonparametric ANOVA (df = 4).

Given these idiosyncratic effects of genetic background, we chose to examine one of the cases in greater detail. In particular, we asked when during its 50,000-generation history did the evolvability of the Ara+5 background decline with respect to tetracycline. To address this question, we examined clones isolated during this population’s early history and identified when the bacteria had lost their capacity to evolve tetracycline resistance during a single exposure, to an extent commensurate with the ancestral strain’s evolvability. As shown in Figure 6, the reduced evolvability was evident in all of the clones isolated from generation 2,000 onward as well as in one of two clones isolated at generation 1,500. With one exception, all of the LTEE-derived parental backgrounds across this time-series had the same MIC value as the ancestor (Fig. 6a). However, the evolved mutants from the later-generation Ara+5 genetic backgrounds had progressively lower levels of tetracycline resistance (Fig. 6b), which when coupled with the same initial resistance level indicates they had become less evolvable in this respect (Fig. 6c). A Kruskal-Wallis test decisively rejects the null hypothesis of equal evolvabilities across the entire set of clones (χ^2^ = 67.89, df = 12, *p* < 0.0001), and Dunnett’s tests comparing the evolvability of each derived clone from LTEE population Ara+5 with that of the ancestor support the temporal breakpoint described above (Table 5).

**Table 5.** Statistical analyses comparing tetracycline resistance evolvability of clones isolated from the Ara+5 population at different generations to the LTEE ancestor.

| Strain | Difference | Lower 95% CI | Upper 95% CI | *p* |
| --- | --- | --- | --- | --- |
| 0.5A | -0.3 | -0.9 | 0.3 | 0.6921 |
| 0.5B | -0.3 | -0.9 | 0.3 | 0.6921 |
| 1A | 0.0 | -0.6 | 0.6 | 1 |
| 1B | -0.2 | -0.8 | 0.4 | 0.9607 |
| 1.5B | -0.3 | -0.9 | 0.3 | 0.6921 |
| 1.5A | -0.6 | -1.2 | 0.0 | 0.0412 |
| 2A | -1.0 | -1.6 | -0.4 | < 0.0001 |
| 2B | -0.8 | -1.4 | -0.2 | 0.0021 |
| 5A | -0.7 | -1.3 | -0.1 | 0.0101 |
| 5B | -1.2 | -1.8 | -0.6 | < 0.0001 |
| 10A | -1.2 | -1.8 | -0.6 | < 0.0001 |
| 10B | -1.3 | -1.9 | -0.7 | < 0.0001 |

Analyses were performed using a Dunnett's test. Strain identifiers begin with a number that corresponds to the generation (in thousands) of their isolation, followed by an arbitrary letter; strains 2A and 2B, for example, are two clones isolated at generation 2,000 of the LTEE.

**Figure 6.**
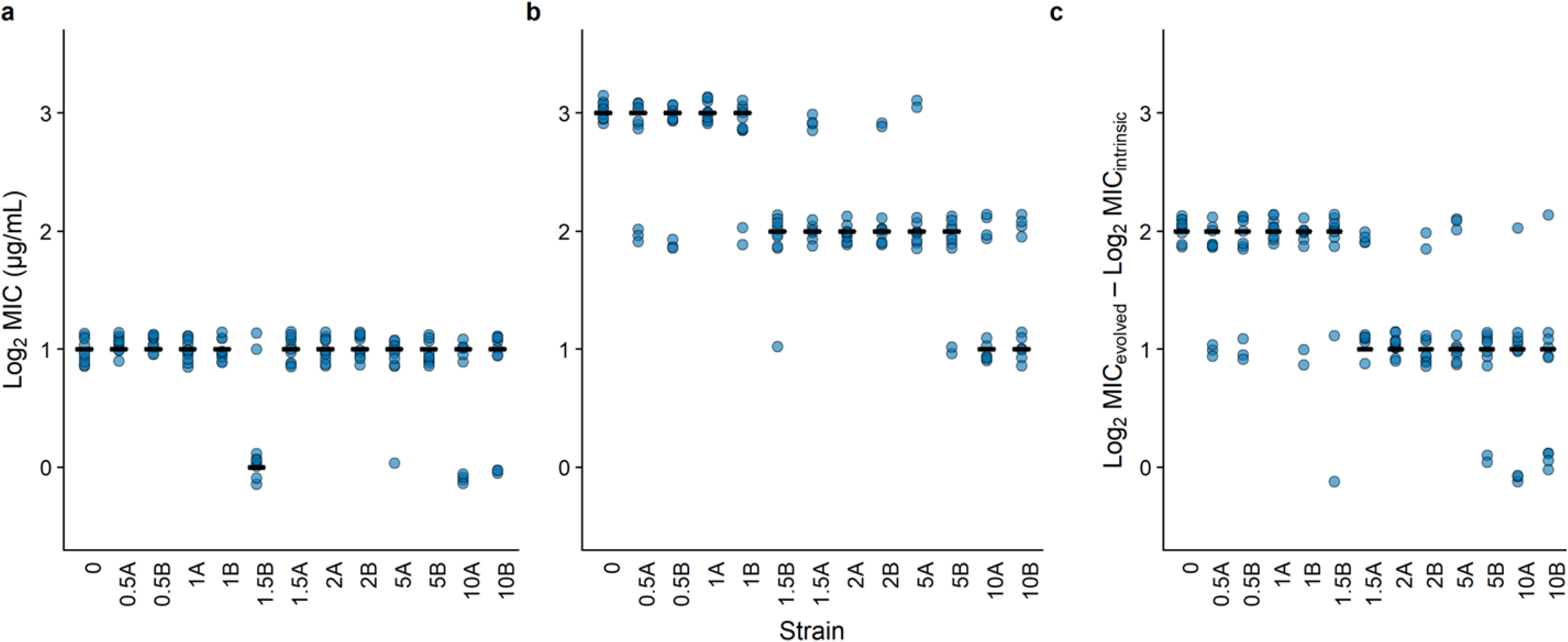
Capacity to evolve resistance to tetracycline in a single step was diminished early in one LTEE lineage. (**a**) Comparison of the intrinsic tetracycline resistance of the ancestor and a time-series of derived strains isolated from the Ara+5 population. Strains are ordered by their time of isolation. The strain identifiers begin with a number corresponding to the generation (in thousands) of their isolation, followed by an arbitrary letter; strains 2A and 2B, for example, are two clones isolated at generation 2,000 of the LTEE. (**b**) Comparison of the evolved resistance levels after one round of drug selection, based on the MICs of the mutant daughter cells derived from the corresponding parental strains. MICs are shown on a log_2_-transformed scale to reflect the fact that the concentrations of antibiotics were tested across a series of two-fold dilutions. (**c**) Evolvability is quantified for each strain as the difference in the log_2_-transformed MICs of the parental strain and its corresponding daughter mutant. Points show 10 independent replicates per strain. Horizontal bars show the median log_2_-transformed MICs (**a, b**) for 10 replicate assays and the evolvability (**c**) based on the corresponding 10 paired differences.

## Discussion

In this study, we addressed a fundamental question about how relaxed selection on a particular set of organismal traits affects their evolvability in situations where those traits again become advantageous. The traits we studied are resistances to several antibiotics, and the question of how changes in genetic background that occur during relaxed selection affects the subsequent evolvability of resistance has potentially important implications for public health. To address these issues, we examined the capacity of *E. coli* strains to evolve increased resistance to four different antibiotics after they had evolved in a drug-free environment for 50,000 generations as part of the long-term evolution experiment (LTEE).

We confirmed that intrinsic resistance tended to decay among the LTEE-derived clones to all four antibiotics we tested (Fig. 4, Table 2). Our results are consistent with a recent study that examined losses of intrinsic resistance in all 12 LTEE lines at generations 2,000 and 50,000 (Lamrabet et al. 2019). Unlike that previous work, however, we then also examined whether and how the LTEE-derived bacteria had changed in their evolvability, specifically their potential to evolve resistance when challenged across a range of concentrations of the same four antibiotics.

We examined two alternative hypotheses that might bear on resistance evolvability. The first is called diminishing returns (Fig. 1b), and it often characterizes the course of adaptive evolution (Moore et al. 2000; Orr 2005; Barrick et al. 2010; Khan et al. 2011; Wiser et al. 2013; Kryazhimskiy et al. 2014; Passagem-Santos et al. 2018). For example, one study used rifampicin-resistant mutants to examine the relation between their initial fitness costs in the absence of this drug and their ability to reduce or eliminate those costs during subsequent evolution, again in a drug-free environment (Barrick et al. 2010). They first isolated eight *rpoB* mutants after a single round of antibiotic selection and showed that the mutants varied in their fitness defects. The authors then propagated these mutants and detected the first beneficial mutations to sweep to high frequency in those populations. They found that the lower-fitness backgrounds gave rise to mutations that conferred greater advantages than did the backgrounds that initially had higher fitness, in accordance with a diminishing-returns model.

If the evolution of antibiotic resistance after a period of decay under relaxed selection conformed to the diminishing-returns model, then we would expect the more susceptible LTEE-derived backgrounds to be more evolvable than their common ancestor. However, we found little statistical support for diminishing returns in our study (Fig. 5, Table 3). There was one instance in which an individual LTEE-derived clone was significantly more evolvable than the ancestor in the ampicillin environment, and two other clones trended in this direction (Fig. 5a, Table S2). However, the statistical support, even for ampicillin, was marginal at best when the evolvabilities of the four clones were analyzed together to account for multiple tests of the same hypothesis (Table 3). In any case, diminishing returns was not typical across the entire set of experiments.

The absence of an overall trend toward diminishing returns might be attributable, in part, to two methodological issues. First, it might point to a limitation of our plate-based approach, and conventional MIC assays in general, to discern subtle differences in MICs and hence in evolvabilities based on differences in MIC values. That is, slight differences in evolvability may be obscured by the discrete resolution of the assays using two-fold increasing concentrations of an antibiotic. Consistent with this possibility, the range of initial susceptibilities was greatest in the ampicillin environment, where the trend toward diminishing returns was most evident (Fig. 5a). An alternative approach that might better capture subtle trends would be to use a continuous culture device that dynamically adjusts drug concentration in the growth medium to match the on-going adaptive dynamics of the population under study (Toprak et al. 2012). Second, we might have had insufficient statistical power to resolve diminishing-returns trends in evolvability. We tested four LTEE-derived lines and their ancestor, whereas some other studies that show diminishing returns in other contexts have used as many as hundreds of lines (Kryazhimskiy et al. 2014).

There is a third factor—one that is biological, rather than methodological—that could also obscure any tendency toward diminishing returns, and that is idiosyncratic heterogeneity among genetic backgrounds in their evolvability (Fig. 1c). This pattern occurs when particular mutations that arose during relaxed selection happen to either constrain or potentiate a strain’s future evolutionary potential with respect to a given selective pressure. We found that the capacity to evolve ceftriaxone resistance actually tended to be lower for the derived clones than for the ancestor (Fig. 5b, Table 4). That tendency is in striking contrast with the ampicillin environment, considering that both drugs are β-lactams and target cell-wall synthesis. This result suggests that these independently derived backgrounds share one or more mutations (or mutated pathways more broadly) that negatively interact with potential mutations that confer ceftriaxone resistance. In a similar vein, we also demonstrated that the Ara+5 lineage had become significantly constrained in its ability to evolve tetracycline resistance relative to both its ancestor and the other LTEE-derived lineages (Fig. 5d).

We conclude, therefore, that historical contingency has played an important role in the capacity of the LTEE-derived populations to respond evolutionarily to changed environments, in particular when challenged with antibiotics. That is, different lineages accumulated genetic differences—even in replicate populations that evolved in the same environment—that influence their ability to evolve and adapt in new directions. Several other microbial-evolution studies have also documented cases of historically contingent outcomes. For example, the mutations that accumulated in one LTEE population potentiated the subsequent evolution of a novel metabolic capacity that arose in only that one population, despite comparable time and opportunity in the eleven other replicate populations (Blount et al. 2008, Quandt et al. 2015, Leon et al. 2018). Similarly, an experiment with *Pseudomonas aeruginosa* showed that high-level colistin resistance was potentiated by prior mutations in transcriptional regulators *phoQ* and *pmrB* (Jochumsen et al. 2016). However, the consequences of contingency in these two cases are the opposite of what we saw in our study: namely, evolvability was potentiated in these previous studies, whereas it became more constrained in ours.

When did the evolvability with respect to tetracycline exposure decline in the Ara+5 population? And why, in molecular-genetic terms, did it decline? To answer the first question, we tested clones from throughout this population’s history and identified when the bacteria first lost their ability to evolve resistance to the same degree as the ancestral strain. We found that this constraint was already present in one of two clones sampled at 1,500 generations of the LTEE, and it was evident in all of the clones we tested from generation 2,000 and onward (Fig. 6). These data thus confirm the idiosyncratic effects of genetic background on the evolvability of resistance, while also greatly narrowing the number of mutations that could have given rise to this change in evolvability. With respect to the second question, we do not yet know the answer, but the timing allows us to narrow the genetic possibilities. By combining our phenotypic results with previously obtained genomic data (Tenaillon et al. 2016), we have identified three candidate mutations that alone or in combination could explain this reduced evolvability. These mutations arise in the following genes: *mreB*, which encodes a protein involved in cell-wall structuring; *pykF*, which encodes pyruvate kinase that catalyzes the last step of glycolysis; and *trkH*, which encodes a potassium ion transporter. Interestingly, recent studies have discovered that mutations in *trkH* can cause increased susceptibility to tetracycline through changes to the proton-motive force, and this relationship may depend upon the genetic background (Lázár et al. 2013; Apjok et al. 2019). In future work, we hope to make genetic constructs that will allow us to investigate the genetic basis for the low evolutionary potential of this background when exposed to tetracycline. We will also sequence some of the antibiotic-resistant mutants that evolved in our experiments to test whether there are any systematic differences among the various strains in the genetic targets of the resistant mutations, and whether such differences correlate with the history of relaxed selection and concomitant increased susceptibility.

In summary, we have shown that bacterial evolution in the absence of antibiotic exposure can lead not only to increased susceptibility, but also to genetic background-dependent changes in resistance evolvability when cells are exposed to those drugs. The evolution of resistance can thus depend upon previously accumulated mutations in a historically contingent fashion. We therefore conclude that strategic antibiotic use could benefit not only from surveillance of current resistance levels in pathogens, but also analyses of their potential to evolve increased resistance in the future. This latter objective would be important both on the scale of the individual patient, where effective treatment is paramount, and on a community-wide scale, where judicious efforts to control the spread of drug resistance become critical. We hope that our evolvability-based approach and extensions thereof prove useful in achieving these objectives.

## Acknowledgments

We acknowledge support from a HHMI Gilliam Fellowship, a Russell B. DuVall Scholarship, and a Rudolph Hugh Award (to K.J.C.); a grant from the National Science Foundation (DEB-1451740 to R.E.L.); and the BEACON Center for the Study of Evolution in Action (DBI-0939454). We thank Neerja Hajela for assistance in the laboratory and Chris Adami, Frances Downes, Joshua Franklin, Devin Lake, Chris Waters, and Mike Wiser for valuable discussions.

## Author contributions

K.J.C. and R.E.L. conceived and designed the study; K.J.C., T.L., and J.B.G. performed the experiments; K.J.C. and R.E.L. analyzed all data; K.J.C. wrote associated analysis scripts and prepared figures; K.J.C., T.L., and R.E.L. wrote the paper. All authors approved the final version.

---

**Table S1.** Statistical significance for changes in intrinsic resistance of the individual clones sampled at generation 50,000 of the LTEE for the four antibiotic treatments.

| **Antibiotic** | **Clone** | ***p*** |
| --- | --- | --- |
| Ampicillin | Ara-5 | 0.0080 |
|  | Ara-6 | 0.0039 |
|  | Ara+4 | 0.0080 |
|  | Ara+5 | 0.0080 |
| Ceftriaxone | Ara-5 | 0.0031 |
|  | Ara-6 | 0.0031 |
|  | Ara+4 | 0.0039 |
|  | Ara+5 | 0.0039 |
| Ciprofloxacin | Ara-5 | 0.2177 |
|  | Ara-6 | 0.1004 |
|  | Ara+4 | 0.0149 |
|  | Ara+5 | 0.0039 |
| Tetracycline | Ara-5 | 0.0031 |
|  | Ara-6 | 0.0031 |
|  | Ara+4 | 0.0039 |
|  | Ara+5 | 0.0278 |

Analyses were performed based on a trinomial distribution (Bian et al. 2011), which reflects the many ties in these datasets. The reported *p*-values are one-tailed, which reflects the expectation that resistance should decline under relaxed selection in the antibiotic-free LTEE environment.

**Table S2.** Statistical significance for trends of diminishing-returns resistance evolvability of the individual clones sampled at generation 50,000 of the LTEE for the four antibiotic treatments.

| **Antibiotic** | **Clone** | ***p*** |
| --- | --- | --- |
| Ampicillin | Ara-5 | 0.0456 |
|  | Ara-6 | 0.1094 |
|  | Ara+4 | 0.1592 |
|  | Ara+5 | 0.8408 |
| Ceftriaxone | Ara-5 | 0.9481 |
|  | Ara-6 | 0.9969 |
|  | Ara+4 | 0.9969 |
|  | Ara+5 | 0.9722 |
| Ciprofloxacin | Ara-5 | 0.9250 |
|  | Ara-6 | 0.8778 |
|  | Ara+4 | 0.1964 |
|  | Ara+5 | 0.1222 |
| Tetracycline | Ara-5 | 0.2895 |
|  | Ara-6 | 0.1946 |
|  | Ara+4 | 0.9250 |
|  | Ara+5 | 0.9722 |

Analyses were performed based on a trinomial distribution (Bian et al. 2011), which reflects the many ties in these datasets. The reported *p*-values are one-tailed, which reflects the directional expectation implied by diminishing returns.

## References

Agudelo-Romero, P., F. de la Iglesia, and S. F. Elena. 2008. The pleiotropic cost of host-specialization in *Tobacco etch potyvirus*. Infect. Genet. Evol. 8:806–814.

Andersson, D. I., and D. Hughes. 2010. Antibiotic resistance and its cost: is it possible to reverse resistance? Nat. Rev. Microbiol. 8:260–271.

Apjok, G., G. Boross, Á. Nyerges, G. Fekete, V. Lázár, B. Papp, C. Pál, and B. Csörgő. 2019. Limited evolutionary conservation of the phenotypic effects of antibiotic resistance mutations. Mol. Biol. Evol. msz 109.

Balaban, N. Q., S. Helaine, K. Lewis, M. Ackermann, B. Aldridge, D. I. Andersson, M. P. Brynildsen, D. Bumann, A. Camilli, J. J. Collins, C. Dehio, S. Fortune, J.-M. Ghigo, W.-D. Hardt, A. Harms, M. Heinemann, D. T. Hung, U. Jenal, B. R. Levin, J. Michiels, G. Storz, M. W. Tan, T. Tenson, L. Van Melderen, and A. Zinkernagel. 2019. Definitions and guidelines for research on antibiotic persistence. Nat. Rev. Microbiol. 17:441–448.

Barrick, J. E., M. R. Kauth, C. C. Strelioff, and R. E. Lenski. 2010. Escherichia coli rpoB mutants have increased evolvability in proportion to their fitness defects. Mol. Biol. Evol. 27:1338–1347.

Baym, M., T. D. Lieberman, E. D. Kelsic, R. Chait, R. Gross, I. Yelin, and R. Kishony. 2016. Spatiotemporal microbial evolution on antibiotic landscapes. Science. 353:1147–1151.

Bian, G., M. McAleer, and W.-K. Wong. 2011. A trinomial test for paired data when there are many ties. Math. Comput. Simul. 81:1153–1160.

Blount, Z. D., C. Z. Borland, and R. E. Lenski. 2008. Historical contingency and the evolution of a key innovation in an experimental population of *Escherichia coli*. Proc. Natl. Acad. Sci. 105:7899–7906.

Blount, Z. D., R. E. Lenski, and J. B. Losos. 2018. Contingency and determinism in evolution: Replaying life’s tape. Science. 362:eaam5979.

Bouma, J. E., and R. E. Lenski. 1988. Evolution of a bacteria/plasmid association. Nature 335:351–352.

Chao, L., B. R. Levin, and F. M. Stewart. 1977. A complex community in a simple habitat: an experimental study with bacteria and phage. Ecology 58:369–378.

Coffey, L. L., and M. Vignuzzi. 2011. Host alternation of Chikungunya virus increases fitness while restricting population diversity and adaptability to novel selective pressures. J. Virol. 85:1025–1035.

Cooper, V. S., A. F. Bennett, and R. E. Lenski. 2001. Evolution of thermal dependence of growth rate of *Escherichia coli* populations during 20,000 generations in a constant environment. Evolution. 55:889–896.

Cooper, V. S., and R. E. Lenski. 2000. The population genetics of ecological specialization in evolving *Escherichia coli* populations. Nature 407:736–739.

Cox, G., and G. D. Wright. 2013. Intrinsic antibiotic resistance: Mechanisms, origins, challenges and solutions. Int. J. Med. Microbiol. 303:287–292.

Darwin, C. 1859. On the Origin of Species. J. Murray, London.

Duffy, S., P. E. Turner, and C. L. Burch. 2006. Pleiotropic costs of niche expansion in the RNA bacteriophage Φ6. Genetics 172:751–757.

Ellis, C. N., and V. S. Cooper. 2010. Experimental adaptation of *Burkholderia cenocepacia* to onion medium reduces host range. Appl. Environ. Microbiol. 76:2387–2396.

Fisher, R. A. 1934. Statistical Methods for Research Workers. 5th ed. Oliver and Boyd, Edinburgh; London.

Han, F., S. Pu, F. Wang, J. Meng, and B. Ge. 2009. Fitness cost of macrolide resistance in *Campylobacter jejuni*. Int. J. Antimicrob. Agents 34:462–466.

Jochumsen, N., R. L. Marvig, S. Damkiær, R. L. Jensen, W. Paulander, S. Molin, L. Jelsbak, and A. Folkesson. 2016. The evolution of antimicrobial peptide resistance in *Pseudomonas aeruginosa* is shaped by strong epistatic interactions. Nat. Commun. 7:13002

Kassen, R., and T. Bataillon. 2006. Distribution of fitness effects among beneficial mutations before selection in experimental populations of bacteria. Nat. Genet. 38:484–488.

Khan, A. I., D. M. Dinh, D. Schneider, R. E. Lenski, and T. F. Cooper. 2011. Negative epistasis between beneficial mutations in an evolving bacterial population. Science. 332:1193–1196.

Koschwanez, J. H., K. R. Foster, and A. W. Murray. 2013. Improved use of a public good selects for the evolution of undifferentiated multicellularity. eLife 2:e00367.

Kryazhimskiy, S., D. P. Rice, E. R. Jerison, and M. M. Desai. 2014. Global epistasis makes adaptation predictable despite sequence-level stochasticity. Science. 344:1519–1522.

Lahti, D. C., N. A. Johnson, B. C. Ajie, S. P. Otto, A. P. Hendry, D. T. Blumstein, R. G. Coss, K. Donohue, and S. A. Foster. 2009. Relaxed selection in the wild. Trends Ecol. Evol. 24:487–496.

Lamrabet, O., M. Martin, R. E. Lenski, and D. Schneider. 2019. Changes in intrinsic antibiotic susceptibility during a long-term evolution experiment with *Escherichia coli*. mBio 10:e00189–19.

Lázár, V., G. Pal Singh, R. Spohn, I. Nagy, B. Horváth, M. Hrtyan, R. Busa-Fekete, B. Bogos, O. Méhi, B. Csörgő, G. Pósfai, G. Fekete, B. Szappanos, B. Kégl, B. Papp, and C. Pál. 2013. Bacterial evolution of antibiotic hypersensitivity. Mol. Syst. Biol. 9:700.

Leiby, N., and C. J. Marx. 2014. Metabolic erosion primarily through mutation accumulation, and not tradeoffs, drives limited evolution of substrate specificity in *Escherichia coli*. PLoS Biol. 12:e1001789

Lenski, R. E. 1988. Experimental studies of pleiotropy and epistasis in *Escherichia coli*. I. Variation in competitive fitness among mutants resistant to virus T4. Evolution 42:425–432.

Lenski, R. E. 1997. The cost of antibiotic resistance from the perspective of a bacterium. Pp. 131–151 in D. J. Chadwick and J. Goode, eds. Antibiotic Resistance: Origins, Evolution, Selection and Spread. Wiley: Chichester, UK.

Lenski, R. E., M. R. Rose, S. C. Simpson, and S. C. Tadler. 1991. Long-term experimental evolution in *Escherichia coli*. I. Adaptation and divergence during 2,000 generations. Am. Nat. 138:1315–1341.

Lenski, R. E., S. C. Simpson, and T. T. Nguyen. 1994. Genetic analysis of a plasmid-encoded, host genotype-specific enhancement of bacterial fitness. J. Bacteriol. 176:3140–3147.

Leon, D., S. D’Alton, E. M. Quandt, and J. E. Barrick. 2018. Innovation in an *E. coli* evolution experiment is contingent on maintaining adaptive potential until competition subsides. PLoS Genet. 14:e1007348.

Luria, S. E., and M. Delbrück. 1943. Mutations of bacteria from virus sensitivity to virus resistance. Genetics 28:491–511.

Melnyk, A. H., A. Wong, and R. Kassen. 2015. The fitness costs of antibiotic resistance mutations. Evol. Appl. 8:273–283.

Meyer, J. R., D. T. Dobias, S. J. Medina, L. Servilio, A. Gupta, and R. E. Lenski. 2016. Ecological speciation of bacteriophage lambda in allopatry and sympatry. Science. 354:1301–1304.

Moore, F. B.-G., D. E. Rozen, and R. E. Lenski. 2000. Pervasive compensatory adaptation in *Escherichia coli*. Proc. R. Soc. London B 267:515–522.

Nguyen, T. N. M., Q. G. Phan, L. P. Duong, K. P. Bertrand, and R. E. Lenski. 1989. Effects of carriage and expression of the Tn*10* tetracycline resistance operon on the fitness of *Escherichia coli* K12. Mol. Biol. Evol. 6:213–225.

Orr, H. A. 2005. The genetic theory of adaptation: A brief history. Nat. Rev. Genet. 6:119–127.

Palmer, A. C., R. Chait, and R. Kishony. 2018. Nonoptimal gene expression creates latent potential for antibiotic resistance. Mol. Biol. Evol. 35:2669–2684.

Passagem-Santos, D., S. Zacarias, and L. Perfeito. 2018. Power law fitness landscapes and their ability to predict fitness. Heredity 121:482–498.

Quandt, E. M., J. Gollihar, Z. D. Blount, A. D. Ellington, G. Georgiou, and J. E. Barrick. 2015. Fine-tuning citrate synthase flux potentiates and refines metabolic innovation in the Lenski evolution experiment. eLife 4:e09696.

Ratcliff, W. C., R. F. Denison, M. Borrello, and M. Travisano. 2012. Experimental evolution of multicellularity. Proc. Natl. Acad. Sci. 109:1595–1600.

Reboud, X., and G. Bell. 1997. Experimental evolution in *Chlamydomonas*. III. Evolution of specialist and generalist types in environments that vary in space and time. Heredity 78:507–514.

Reynolds, M. G. 2000. Compensatory evolution in rifampicin-resistant *Escherichia coli*. Genetics 156:1471–1481.

Rozen, D. E., L. McGee, B. R. Levin, and K. P. Klugman. 2007. Fitness costs of fluoroquinolone resistance in *Streptococcus pneumoniae*. Antimicrob. Agents Chemother. 51:412–416.

Schrag, S. J., V. Perrot, and B. R. Levin. 1997. Adaptation to the fitness costs of antibiotic resistance in *Escherichia coli*. Proc. R. Soc. B 264:1287–1291.

Sokal, R. R., and F. J. Rohlf. 1994. Biometry, 3rd ed. W. H. Freeman.

Tenaillon, O., J. E. Barrick, N. Ribeck, D. E. Deatherage, J. L. Blanchard, A. Dasgupta, G. C. Wu, S. Wielgoss, S. Cruveiller, C. Médigue, D. Schneider, and R. E. Lenski. 2016. Tempo and mode of genome evolution in a 50,000-generation experiment. Nature 536:165–170.

Toprak, E., A. Veres, J.-B. Michel, R. Chait, D. L. Hartl, and R. Kishony. 2012. Evolutionary paths to antibiotic resistance under dynamically sustained drug selection. Nat. Genet. 44:101–105.

Turner, P. E., and S. F. Elena. 2000. Cost of host radiation in an RNA virus. Genetics 156:1465–1470.

Wasik, B. R., A. Bhushan, C. B. Ogbunugafor, and P. E. Turner. 2015. Delayed transmission selects for increased survival of vesicular stomatitis virus. Evolution. 69:117–125.

Wenger, J. W., J. Piotrowski, S. Nagarajan, K. Chiotti, G. Sherlock, and F. Rosenzweig. 2011. Hunger artists: Yeast adapted to carbon limitation show trade-offs under carbon sufficiency. PLoS Genet. 7:e1002202.

Wiser, M. J., N. Ribeck, and R. E. Lenski. 2013. Long-term dynamics of adaptation in asexual populations. Science. 342:1364–1367.

